# rhythmCI: an R package for rhythm detection using harmonic regression amplitude confidence intervals

**DOI:** 10.64898/2026.10.05.756647

**Authors:** Jayanth Sreekanth, Andreas Edel, Claudia Spies, Sandro Andreotti, Bharath Ananthasubramaniam

## Abstract

**Background:** Circadian clocks are evolutionarily conserved systems that orchestrate daily rhythms in human physiology, tissue behavior, and cellular function. In circadian research, the NanoString nCounter platform has emerged as a robust alternative to microarray and RNA sequencing. However, raw instrument output requires specialized preprocessing to handle technical noise. Furthermore, while harmonic regression is widely used to detect periodic signals in biological time-series data, evaluating rhythmicity remains highly challenging in clinical studies characterized by sparse sampling and small sample sizes.

**Results:** To address these limitations, we developed rhythmCI, an easy-to-use R package that integrates established normalization methods with a complete workflow for circadian data analysis. The package incorporates a bootstrapping procedure to estimate confidence intervals for harmonic regression amplitudes, offering a statistically transparent way to quantify rhythmicity in small-cohort studies. rhythmCI provides an end-to-end workflow for NanoString nCounter data, including automated preprocessing, normalization, and visualization. We evaluated the utility and accuracy of the package using synthetic time-series data generated with the Simphony framework, demonstrating that rhythmCI produces interpretable and reliable rhythmicity classifications under controlled simulation conditions.

**Conclusions:** rhythmCI bridges a critical gap in circadian bioinformatics by offering a reliable method to characterize periodicity under challenging experimental designs typical of clinical research. By streamlining NanoString workflows and improving the statistical rigor of rhythm detection in small datasets, rhythmCI enhances the accessibility and reliability of circadian biology analysis.

## Background

Circadian clocks are evolutionarily conserved, endogenous timing systems driven by cell-autonomous transcriptional-translational feedback loops involving core clock components such as Period genes (PER1, PER2, PER3), Cryptochrome 1 (CRY1), and Aryl Hydrocarbon Receptor Nuclear Translocator Like (ARNTL, also known as BMAL1)[1]. These molecular oscillators coordinate physiology and behavior by aligning internal rhythms with external light–dark cycles through robust environmental entrainment[2]. However, in settings like the Intensive Care Unit (ICU), the absence of natural light combined with severe physiological stressors, such as sepsis, acute inflammation, and metabolic stress, profoundly disrupts clock entrainment[3]. Consequently, studies report dampened circadian rhythms in serum melatonin and core clock gene expression among critically ill patients[4-6]. Yet, because dense time-series blood sampling is rarely feasible in such an environment, the scientific literature on clock-gene regulation during critical illness remains limited.

The NanoString nCounter platform has become increasingly popular for quantifying mRNA expression in clinical studies[7]. Unlike microarrays or RNA-Seq, NanoString uses digital color-coded barcodes to directly measure up to ∼800 transcripts without enzymatic amplification, eliminating amplification bias and enabling targeted, cost-efficient profiling[8]. As the nCounter platform is increasingly used in longitudinal clinical studies, technical variation must be removed efficiently without compromising underlying biological signals. Robust preprocessing is therefore essential, and the “Remove Unwanted Variation with negative control genes” (RUVg)-based normalization algorithm[9] has been shown to outperform the traditional nSolver workflow[10] in preserving biological variation while reducing technical artifacts [11], establishing it as a widely used and important method for accurate rhythm discovery in clinical cohorts.

Characterizing gene expression oscillations from hospitalized patients is central to modern circadian medicine, yet standard analytical approaches struggle under real-world sampling constraints. Ideal circadian studies collect samples every 1–4 hours[12], but clinical datasets are typically sparse, irregularly spaced, or truncated, leaving traditional nonparametric methods statistically underpowered[13]. Parametric strategies, such as harmonic regression, offer greater sensitivity by explicitly modeling waveform parameters like amplitude and acrophase[12], enabling detection of weak or low-amplitude rhythms even in small cohorts[14].

To address the dual challenges of NanoString-specific technical noise and the sparse sampling typical of clinical cohorts, we created rhythmCI, a unified framework that enables efficient preprocessing and normalization of NanoString expression data and implements rhythmicity quantification using the approach previously developed by Lachmann et al.[6] through bootstrapped amplitude confidence intervals (CIs). This regression-based method for quantifying rhythmicity is broadly applicable, not only to gene-expression datasets, but also to other biological time-series measurements such as protein expression. rhythmCI pairs this generalizable rhythmicity analysis with informative visualizations that accompany each result.

## Implementation

### Installation

rhythmCI is an open-source R package available through GitHub, and can be installed using standard tools such as pak[15] or remotes[16]. By minimizing external dependencies, rhythmCI enables seamless integration into existing analytical workflows for circadian rhythm detection.

### Usage

The rhythmCI package provides a practical, end-to-end workflow for circadian rhythm analysis in NanoString and other biological time-series datasets. Users can load raw or pre-processed expression data, apply RUVg-based normalization[9], fit harmonic regression models, estimate bootstrapped amplitude CIs, and generate publication-ready visualizations.

rhythmCI accepts multiple input formats, including Reporter Code Count (RCC) files, count matrices, and processed data frames, and supports batch processing across many genes or targets. The package outputs amplitude, CIs for amplitude, the Midline Estimating Statistic of Rhythm (MESOR) and acrophase, and provides exportable diagnostic plots for downstream analysis.

A detailed vignette is included with the installation and available online, demonstrating each step of the workflow with examples.

### Inputs

The rhythmCI workflow accepts two types of input depending on whether NanoString preprocessing has already been performed.

1. RCC directory input: Users who wish to load and normalize NanoString nCounter data in a single step can supply a directory containing RCC files along with a separately defined metadata file. In this metadata file, the first column must list RCC file names, and subsequent columns should specify sample label, sample time, and any additional relevant characteristics. A template metadata file (*meta_GSE146530*.*csv*) is included with the package to guide users in preparing and adjusting their own metadata. Any duplicated entries in the metadata table are automatically removed, retaining only the first occurrence to ensure consistent sample-to-file mapping.
2. Pre-processed table input: If quality control has already been completed and the NanoString data are available in a tabular format, this table can be provided directly. Users can then proceed with normalization or rhythm detection without needing to load RCC files or a metadata sheet.

These two input pathways allow rhythmCI to flexibly integrate into both raw-data workflows and established preprocessing pipelines.

### Normalization

rhythmCI provides three normalization options: “none”, “nSolver[10]”, and “RUVg[9]”. The appropriate choice depends on the quality of the NanoString data and whether genes fall below the Limit of Detection (LOD). An overview of steps is shown in Figure 1.

**Figure 1.**
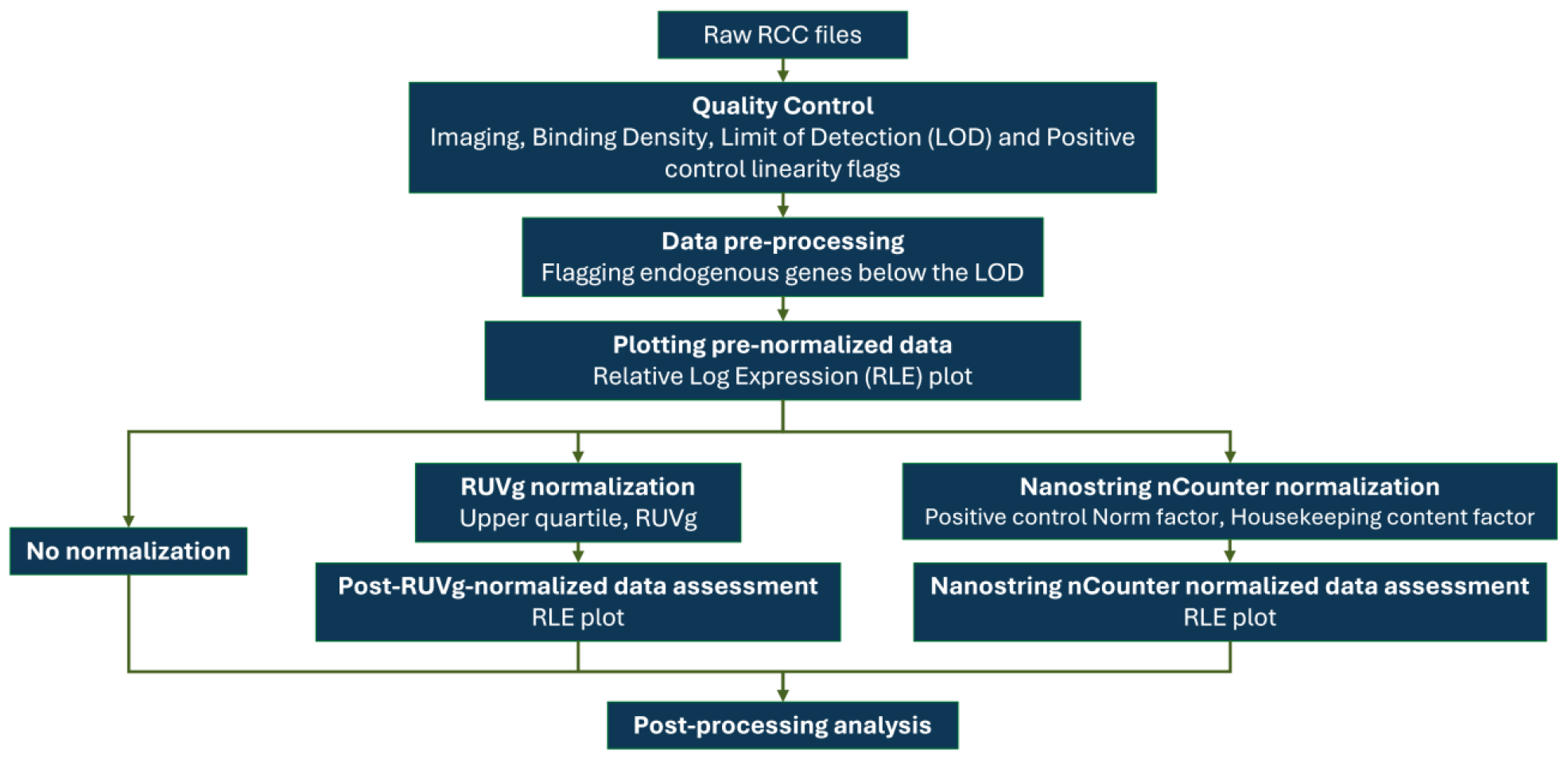
Overview of the normalization workflow applied when raw RCC files are provided, illustrating the sequential steps from data import through quality control and normalization.

Abbreviations: RCC, Reporter Code Count; LOD, Limit of Detection; RLE, Relative Log Expression; RUVg, Remove Unwanted Variation with negative control genes

When performing initial quality control, we recommend selecting “none”. This allows users to inspect LOD metrics and follow the guidelines to identify and flag problematic genes. Any genes failing quality thresholds can then be handled according to user-specific criteria.

If one or more genes are filtered during quality control, the dataset can be normalized using the RUVg-based method[9] via the *normalizeruvg* function. If no genes fall below quality metrics, users may choose either RUVg or nSolver normalization through the *loadnormalizercc* function.

Because RUVg-based normalization relies on housekeeping genes, users must ensure that these genes are stable and not associated with the biology under investigation. We recommend the article by Gagnon et al.[17] for guidance on selecting an appropriate k-value.

After normalization, users can generate Relative Log Expression (RLE) plots[18] from binary logarithm (log2)-transformed expression or normalized data to assess sample-level consistency.

### Detecting rhythmicity

A widely used parametric approach for detecting rhythmicity in biological time-series data is harmonic regression, in which a cosine-based regression model is fit to the observations[19]. Using least-squares estimation, the model describes rhythmic patterns with three parameters: acrophase (the time of peak expression), amplitude (half the peak-to-trough difference), and MESOR (the mean level of the time series). The period is fixed and selected according to the organism or experimental design.

The model is expressed as

**Equation 1**

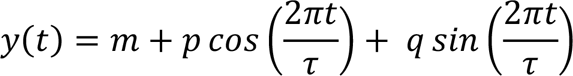

where *y*(*t*) represents gene expression/parameter value at time *t, m* is the MESOR, and τ is the period. The parameters amplitude (*A*) and acrophase (Φ) can be derived from the coefficients *p* and *q* as follows:

**Equation 2**

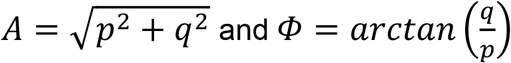

To perform harmonic regression, rhythmCI requires a matrix with samples across time in rows and variables (e.g., genes) in columns. After fitting the cosine model, the best-fit curve and residuals are obtained. Residuals are then resampled with replacement using bootstrapping. For each bootstrap iteration, sampled residuals are added to the best-fit curve to generate a bootstrap estimate of amplitude. CIs for amplitude are computed using the bias-corrected and accelerated method[20, 21]. The user can specify the alpha level, with optional Bonferroni correction.

The MESOR, amplitude, and acrophase are estimated before bootstrapping. Using the CI of amplitude and a user-defined *fc_cutoff*, rhythmCI assigns rhythmic status to each variable. Because *fc_cutoff* represents the log2 fold amplitude change, a value of 0.5 (1.4 on the normal scale) is generally considered biologically meaningful[12]. Based on this cutoff, each variable is classified as:

- Rhythmic — both CI bounds lie above the cutoff
- Arrhythmic — both CI bounds lie below or equal to the cutoff
- Inconclusive — CI bounds straddle the cutoff

At least four samples per subject are required for rhythmicity analysis. Variables with fewer than four observations will be excluded. When missing values are present, or subjects have unequal numbers of samples, column-wise operations are to be defined, which may require more computation time than matrix-wise processing.

## Results

To demonstrate the functionality of rhythmCI, we applied the workflow to both publicly available NanoString data and synthetic time-series datasets. Normalization is illustrated using the GSE146530 dataset[22]. Examination of sample-wise median deviation in log2 expression through RLE plots (Figure 2) showed that RUVg-normalized data exhibited better sample uniformity compared with nSolver-normalized or raw expression values, highlighting the effectiveness of RUVg in reducing technical variation.

**Figure 2.**
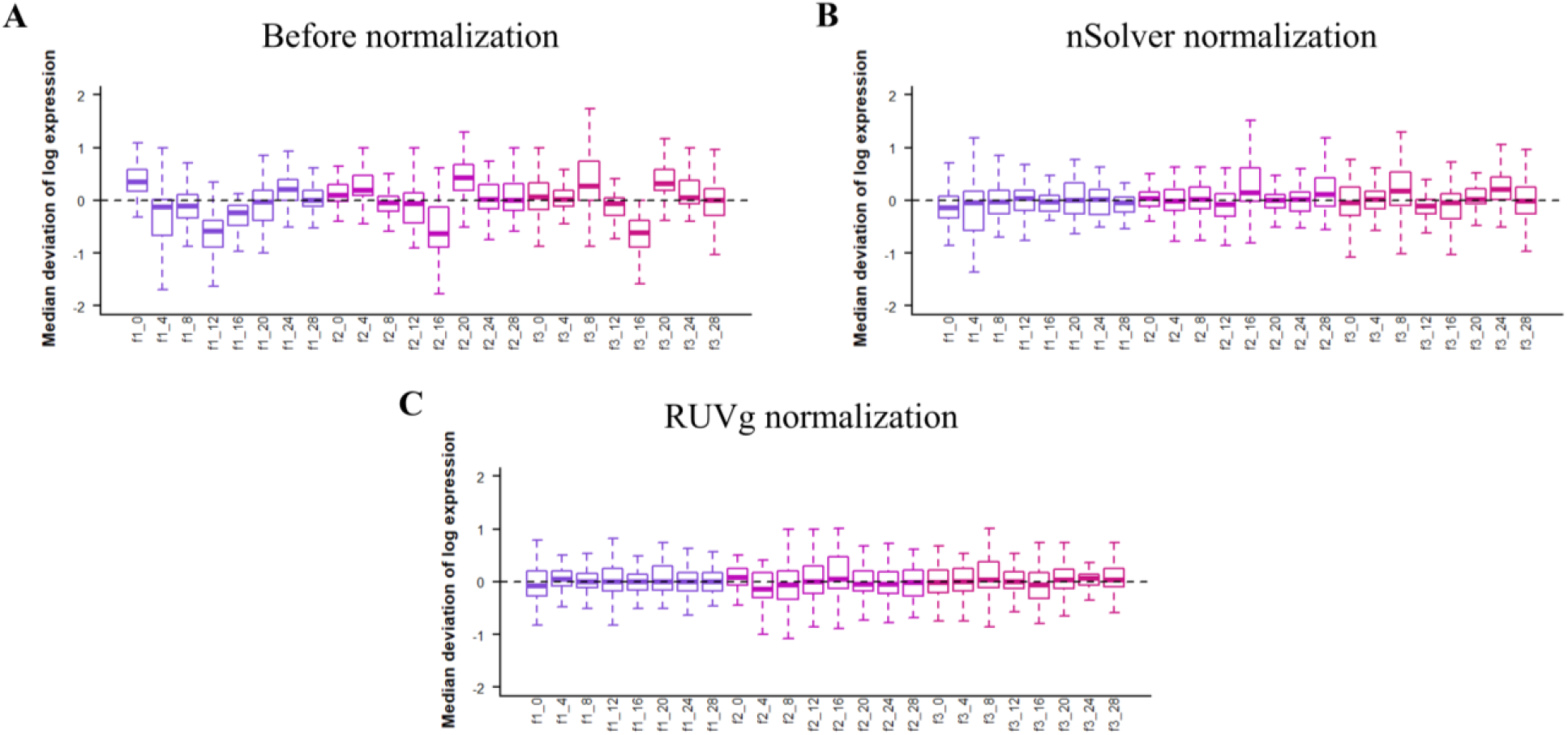
RLE plots showing (A) log2-transformed raw expression values from the GSE146530 dataset before normalization, (B) log2 values after nSolver normalization, and (C) log2 values after RUVg normalization. The x-axis represents samples collected at different time-stamps, and the y-axis shows the relative log expression, calculated as each gene’s deviation from its median log2-transformed expression across all samples. Abbreviations: RLE, Relative Log Expression; RUVg, Remove Unwanted Variation with negative control genes; log2, Binary logarithm

Quantitative evaluation of pre- and post-normalization data including both sample-wise and dataset-wide summary statistics alongside metric definitions is provided in Supplementary File 1. The Complete R script required to compute these metrics is provided in Supplementary File 2.

Following preprocessing and normalization, rhythmicity detection was evaluated using synthetic data generated with Simphony[23], which simulates gene expression from a negative-binomial distribution. We generated expression profiles for 10 genes across 5 patients, each with 7 time-points spanning 0 to 24 hours. The first five genes were assigned low amplitudes (0.1 – 0.49), while the remaining five were assigned higher amplitudes (0.5 – 2.0) on the log2 scale, with low noise dispersion (reduced to 5% of the default variance) specified during simulation. Rhythmicity classification was performed using the *rhythmicity()* function with *operation=“matrix”, seed=7, tau=24, num_boot=3999, alpha=0*.*05, and fc_cutoff=0*.*5*. rhythmCI correctly identified most rhythmic and arrhythmic genes, demonstrating reliable classification performance. Of the 35 total features, 6 arrhythmic genes were labeled inconclusive, and 6 rhythmic genes were also deemed inconclusive. These results indicate that rhythmCI effectively distinguishes rhythmic from non-rhythmic patterns while providing interpretable confidence-interval-based classifications. These CIs provide a way to contrast rhythms in a gene across subjects or conditions and provide a complementary approach to comparing rhythms[24]. The simulated oscillatory expression patterns classified according to their rhythmic categories are shown in Figure 3, and the bootstrap-derived amplitude CIs are presented in Figure 4.

Abbreviations: log2, Binary logarithm

**Figure 3.**
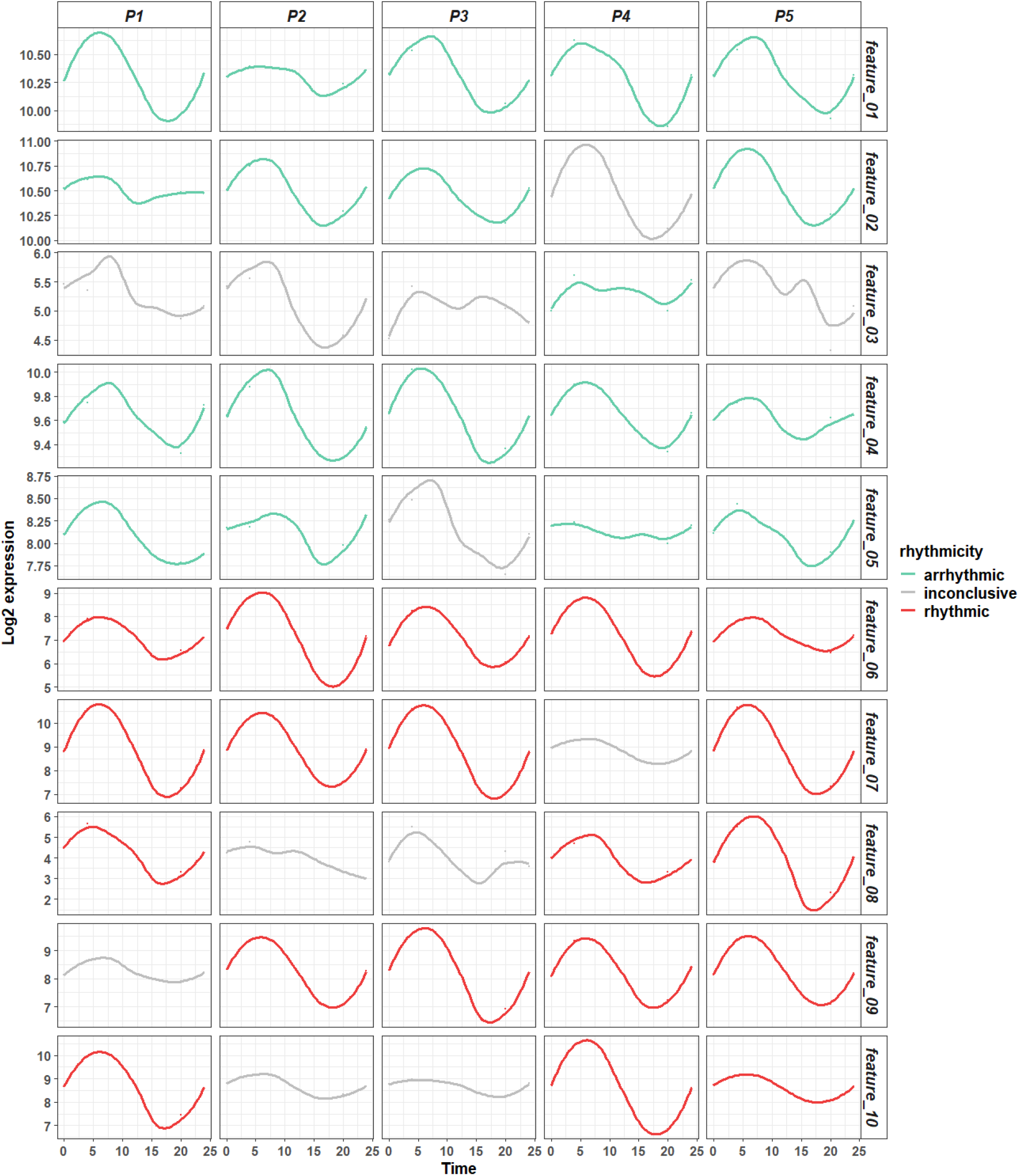
Simulated gene expression patterns across time and the rhythmicity categories (aquamarine3: arrhythmic, grey: inconclusive, brown2: rhythmic) assigned by rhythmCI. The image is faceted such that each row corresponds to a gene (labeled as features) and each column corresponds to a patient (P1-P5). Within each facet, the x-axis represents the sampling time, and the y-axis shows the log2-transformed expression values for that gene.

**Figure 4.**
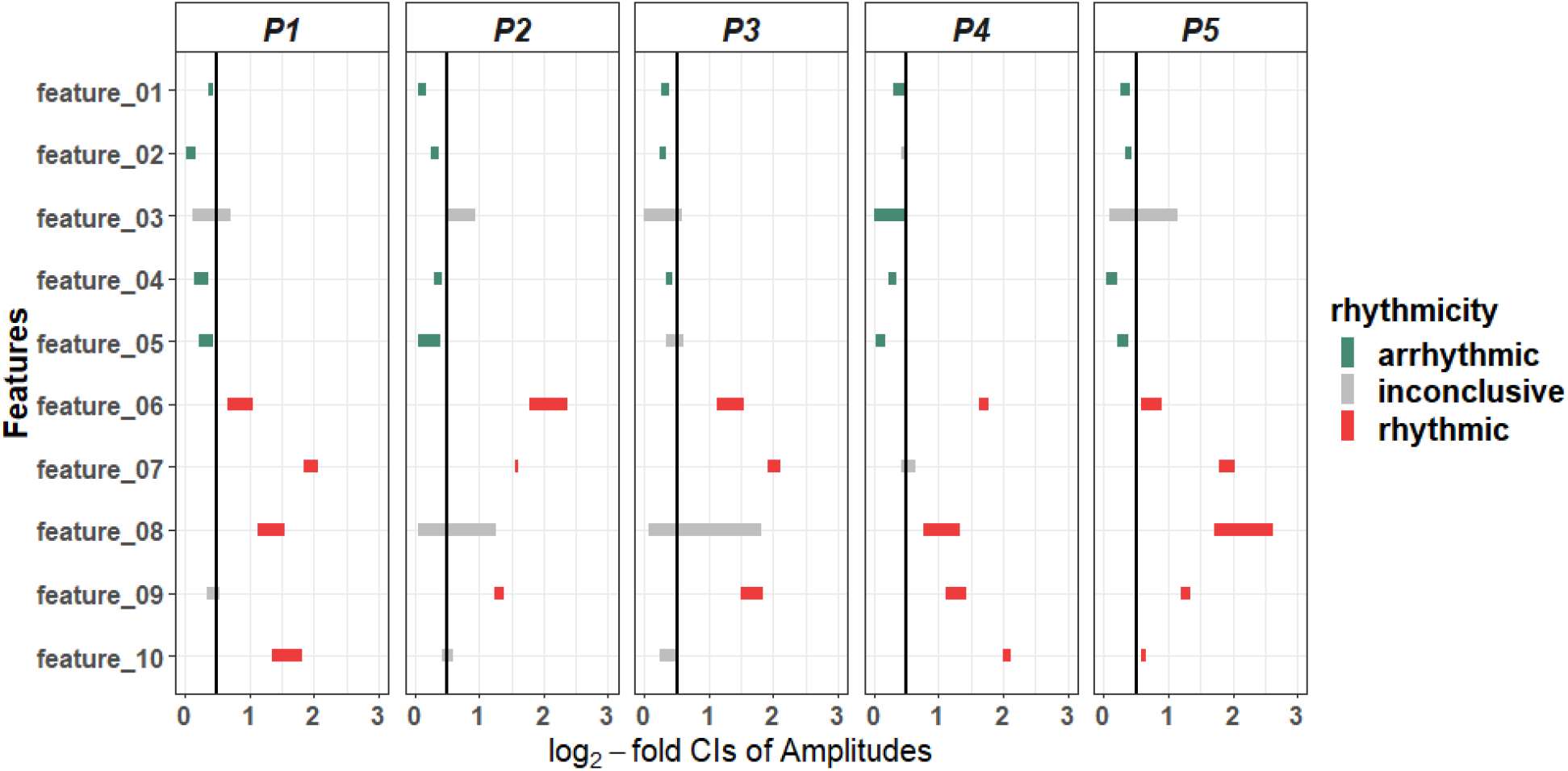
Bootstrap-derived amplitude CIs for each feature, as estimated by rhythmCI, illustrating the confidence-based classification of arrhythmic (aquamarine3), inconclusive (grey), and rhythmic (brown2) genes. The image is faceted patient-wise, with each column corresponding to a patient (P1-P5). Within each facet, the x-axis represents the log2-fold confidence intervals of amplitude estimates, and the y-axis lists genes (labeled as features).

Abbreviations: CI, Confidence Interval

## Conclusions

rhythmCI provides a streamlined and reproducible framework for circadian rhythm analysis in NanoString and other biological time-series datasets. By integrating flexible input handling, established normalization approaches, harmonic regression modeling, and confidence-interval-based classification, the package supports a complete workflow from raw RCC files to interpretable rhythmicity results. The normalization demonstration using the GSE146530 dataset and the simulation dataset generated with Simphony illustrate how rhythmCI can be applied in practice and how its rhythmicity module performs under controlled conditions. As interest in temporal gene regulation continues to expand across experimental and clinical domains, rhythmCI offers a practical and accessible tool for quantifying rhythmic patterns.

## Availability and requirements

Project name: rhythmCI

Project home page: https://github.com/Jayanth-Sreekanth/rhythmCI

Operating system(s): Platform independent

Programming language: R

Other requirements: None

License: GNU GPL v3

Any restrictions to use by non-academics: No

### List of abbreviations

RNA: Ribonucleic acid
MESOR: Midline Estimating Statistic of Rhythm
PER1, PER2, PER3: Period genes
CRY1: Cryptochrome 1
ARNTL: Aryl Hydrocarbon Receptor Nuclear Translocator Like
BMAL1: Basic helix-loop-helix ARNT-like protein 1
ICU: Intensive Care Unit
RNA-Seq: Ribonucleic acid sequencing
mRNA: messenger ribonucleic acid
RUVg: Remove Unwanted Variation with negative control genes
RCC: Reporter Code Count
LOD: Limit of Detection
RLE: Relative Log Expression
log2: Binary logarithm
CI: Confidence Interval

## Declarations

### Ethics approval and consent to participate

Not applicable.

### Consent for publication

Not applicable.

### Availability of data and materials

rhythmCI is open-source and freely accessible as an R package on GitHub: https://github.com/Jayanth-Sreekanth/rhythmCI with example datasets and vignettes included to support reproducible analysis. All materials required to run the functions demonstrated in this manuscript are available through the package repository.

### Competing interests

J.S. reports no conflict of interest with respect to the study. A.E. declares that his institution received payments or honoraria for lectures from Dräger Medical Deutschland GmbH. He holds shares of Novavax, BioNTech and Bavarian (<5000€). C.S. reports grants from the German Research Society (Deutsche Forschungsgemeinschaft), Einstein Foundation Berlin, German Federal Joint Committee (Gemeinsamer Bundesausschuss G-BA) Innovationsfond, Inner University Grants, Georg Thieme Verlag, Dr. F. Köhler Chemie GmbH, Sintetica GmbH, Stifterverband für die Deutsche Wissenschaft e.V. (Medtronic and Teladoc Health), German Federal Ministry of Research, Technology and Space (BMFTR), Robert Koch Institute-a Federal Institute within the portfolio of the German Federal Ministry of Health, the European Commission Horizon Europe, AbbVie Deutschland GmbH & Co. KG, AOP Orphan Pharmaceuticals Germany GmbH (Member of the AOP Health Group), CytoSorbents Europe GmbH, Dr. Franz Köhler Chemie GmbH, Dräger, Löwenstein Medical SE & Co. KG, Medtronic GmbH, Mundipharma Deutschland GmbH & Co. KG, CSL Vifor Pharma Deutschland GmbH, and PAION Pharma GmbH during the conduct of the study. In addition, C.S. has multiple different patents and an unpaid leadership or fiduciary role in the German Research Foundation (Deutsche Forschungsgemeinschaft), German National Academy of Sciences–Leopoldina (Deutsche Akademie der Naturforscher Leopoldina e. V.), Berlin Medical Society (Berliner Medizinische Gesellschaft), ESICM (European Society of Intensive Care Medicine), ESAIC (European Society of Anaesthesiology and Intensive Care), German Society of Anaesthesiology and Intensive Care Medicine (DGAI), German Interdisciplinary Association for Intensive Care and Emergency Medicine (DIVI) and German Sepsis Foundation (Deutsche Sepsis-Stiftung). C.S. has participated in the Data Safety Monitoring Board or Advisory Board of Takeda Pharmaceutical Company Limited, Mundipharma, PAION Pharma GmbH, and Lynx Health Science GmbH. S.A. reports no conflict of interest with respect to the study. B.A. reports grants from the German Research Society (Deutsche Forschungsgemeinschaft). He reports support from RIKEN Wako, Japan for a guest stay in Japan for 10 days in October 2025.

## Funding

B.A. acknowledges support from grants 444137814, 511886499 and 528064681 from Deutsche Forschungsgemeinschaft (DFG). This work was funded by the Deutsche Forschungsgemeinschaft (DFG, German Research Foundation) – Project-ID 541063275 – TRR 418.

The funders had no role in the design of the study, collection, analysis and interpretation of the data and in the preparation of the manuscript.

## Authors’ contributions

C.S. originated the study concept and provided financial support. B.A. developed the core methodology, and J.S. conceived supporting analytical steps to conceptualize the project. B.A. and S.A. supervised the study. J.S. led software package development, data generation, and data analysis. J.S., B.A., S.A., A.E. and C.S. interpreted the results. J.S. wrote the original draft, which was critically reviewed by all authors. All authors read and approved the final manuscript.

## Acknowledgements

The authors thank the contributors of the open-source software packages used in this work. The authors used Google Gemini 3.6 Flash and Microsoft 365 Copilot (GPT-5 model) for writing and limited coding assistance. The primary codebase was developed independently of large language models. All final content and code were reviewed and revised by the authors, who accept complete accountability for the published work.

## Supplementary Files

**supplementary File 1** Quantitative summary of pre- and post-normalization data. Contains sample-wise metrics (medians, IQRs, MADs) and overall dataset-wide summary statistics, along with formal metric definitions.

**supplementary File 2** The text file (Summary_pre_and_post_normalization.txt) containing the R script to reproduce Supplementary File 1.

## Notes

### Summary of Updates

The author affiliations were previously incomplete; full affiliation details have now been provided.

## References

1. Jones SE, Lane JM, Wood AR, Van Hees VT, Tyrrell J, Beaumont RN, Jeffries AR, Dashti HS, Hillsdon M, Ruth KS et al: Genome-wide association analyses of chronotype in 697,828 individuals provides insights into circadian rhythms. Nature Communications 2019, 10(1).

2. Hirota T, Fukada Y: Resetting mechanism of central and peripheral circadian clocks in mammals. Zoolog Sci 2004, 21(4):359–368.

3. Felten M, Dame C, Lachmann G, Spies C, Rubarth K, Balzer F, Kramer A, Witzenrath M: Circadian rhythm disruption in critically ill patients. Acta Physiologica 2023, 238(1).

4. Maas MB, Iwanaszko M, Lizza BD, Reid KJ, Braun RI, Zee PC: Circadian Gene Expression Rhythms During Critical Illness. Crit Care Med 2020, 48(12):e1294–e1299.

5. Diaz E, Diaz I, Del Busto C, Escudero D, Pérez S: Clock Genes Disruption in the Intensive Care Unit. J Intensive Care Med 2020, 35(12):1497–1504.

6. Lachmann G, Ananthasubramaniam B, Wünsch VA, Scherfig L-M, Von Haefen C, Knaak C, Edel A, Ehlen L, Koller B, Goldmann A et al: Circadian rhythms in septic shock patients. Annals of Intensive Care 2021, 11(1).

7. Eastel JM, Lam KW, Lee NL, Lok WY, Tsang AHF, Pei XM, Chan AKC, Cho WCS, Wong SCC: Application of NanoString technologies in companion diagnostic development. Expert Review of Molecular Diagnostics 2019, 19(7):591–598.

8. Tsang H-F, Xue VW, Koh S-P, Chiu Y-M, Ng LP-W, Wong S-CC: NanoString, a novel digital color-coded barcode technology: current and future applications in molecular diagnostics. Expert Review of Molecular Diagnostics 2017, 17(1):95–103.

9. Risso D, Ngai J, Speed TP, Dudoit S: Normalization of RNA-seq data using factor analysis of control genes or samples. Nature Biotechnology 2014, 32(9):896–902.

10. Aboyoun P, Ortogero N, Yang Z, Reeves J, Gorman K, Vitancol R, Smith T, Ren Y, Henderson D: NanoStringNCTools: NanoString nCounter Tools. NanoString Technologies: Seattle, WA, USA 2023.

11. Bhattacharya A, Hamilton AM, Furberg H, Pietzak E, Purdue MP, Troester MA, Hoadley KA, Love MI: An approach for normalization and quality control for NanoString RNA expression data. Briefings in bioinformatics 2021, 22(3):bbaa163.

12. Thaben PF, Westermark PO: Differential rhythmicity: detecting altered rhythmicity in biological data. Bioinformatics 2016, 32(18):2800–2808.

13. Cunningham PS, Kitchen GB, Jackson C, Papachristos S, Springthorpe T, Van Dellen D, Gibbs J, Felton TW, Wilson AJ, Bannard-Smith J et al: ClinCirc identifies alterations of the circadian peripheral oscillator in critical care patients. Journal of Clinical Investigation 2023, 133(4).

14. Baum L, Johns M, Poikela M, Möller R, Ananthasubramaniam B, Prasser F: Data integration and analysis for circadian medicine. Acta Physiologica 2023, 237(4).

15. Csárdi G, Hester J: pak: Another Approach to Package Installation. R package version 0111 2026.

16. Csárdi G, Hester J, Wickham H, Chang W, Morgan M, Tenenbaum D: remotes: R package installation from remote repositories, including ‘GitHub’. R package version 250 2024.

17. Gagnon-Bartsch JA, Speed TP: Using control genes to correct for unwanted variation in microarray data. Biostatistics 2012, 13(3):539–552.

18. Gandolfo LC, Speed TP: RLE plots: Visualizing unwanted variation in high dimensional data. PLOS ONE 2018, 13(2):e0191629.

19. Cornelissen G: Cosinor-based rhythmometry. Theoretical Biology and Medical Modelling 2014, 11(1):16.

20. Puth M-T, Neuhäuser M, Ruxton GD: On the variety of methods for calculating confidence intervals by bootstrapping. Journal of Animal Ecology 2015, 84(4):892–897.

21. Jung K, Lee J, Gupta V, Cho G: Comparison of Bootstrap Confidence Interval Methods for GSCA Using a Monte Carlo Simulation. Frontiers in Psychology 2019, 10.

22. West AC, Iversen M, Jørgensen EH, Sandve SR, Hazlerigg DG, Wood SH: Diversified regulation of circadian clock gene expression following whole genome duplication. PLOS Genetics 2020, 16(10):e1009097.

23. Singer JM, Fu DY, Hughey JJ: Simphony: simulating large-scale, rhythmic data. PeerJ 2019, 7:e6985.

24. Pelikan A, Herzel H, Kramer A, Ananthasubramaniam B: Venn diagram analysis overestimates the extent of circadian rhythm reprogramming. The FEBS Journal 2022, 289(21):6605–6621.

